# Transient dynamics and nonlinear fitness: a unified matrix approach to press and pulse perturbation

**DOI:** 10.1101/2023.10.20.563360

**Authors:** Harman Jaggi, Shripad Tuljapurkar, Wenyun Zuo, Samuel J. L. Gascoigne, Maja Kajin, Roberto Salguero-Gómez

## Abstract

Disturbances can occur as short-lived pulses (e.g., storms) or sustained presses (e.g., chronic drought). Much work in ecology has developed methods to help predict how natural populations respond to disturbances, but analyses of pulse and press disturbances has been largely disconnected. We present a unified matrix framework that links presses and pulses within the same analytical approach, showing how transient nonlinearities and demography shape fitness. We find that transient responses to pulse disturbances accumulate to determine the long-term response to press disturbances. For structured-population models, this cumulative change is given by a new Transient Response Matrix (TRM). Strikingly, the TRM also yields the second derivatives of population growth rate with respect to matrix elements. Thus, there is an intimate but unexpected relationship between nonlinear selection pressures on demographic rates, and the transient dynamics of populations. This relationship yields a strong correlation between TRM and generation time across 439 unique plant and animal species (2690 population models). We also show that the TRM is directly related to Cohen’s cumulative distance measure for populations converging to stability. Our framework provides ecologists with a general tool to predict population responses to diverse environmental changes.

## Introduction

We live in a period of environmental disturbances that may push populations away from their historical states (Barnosky et al. 2012; Abbott et al. 2024). An important goal in Ecology is to predict how natural populations respond to such changes (Morozov et al. 2024). Bender et al. (1984) introduced two terms (pulse and press) to distinguish between two types of experimental disturbances and to describe the combination of cause and effect (Glasby and Underwood 1996). They classified disturbance regimes into acute, discrete events (pulse disturbances) and diffuse, sustained events (press disturbances).

A pulse disturbance is a one-off event (such as an acute epidemic, or extreme but short weather event like a fire or hurricane) that may perturb the system away from its stable state if it fails to resist the disturbance. After a pulse, transient dynamics occur as the system returns to the previous stable state (White et al. 2013; Tao et al. 2021). Examples of a pulse disturbance include preferential targeting of prime adult individuals (Traill et al. 2014) in trophy hunting, droughts that primarily affect juveniles in plant populations (Refsland and Fraterrigo 2018), among others. In contrast a press disturbance is long-lasting (e.g. global warming, land use change) and alters the stable state itself (Donohue et al. 2016; Inamine et al. 2022). In a press, there are also transient dynamics but the repeated disturbances push the population structure away from the original stable state distribution (hereafter, SSD) to the new stable state. Examples of a press perturbation include the decline in fertility rates of a population following chronic disease (Sironi 2019), or the effects of reintroduction of a species into an ecosystem (Ripple and Beschta 2012).

Much attention has been paid theoretically (Yang et al. 2008; Jentsch and White 2019; Hastings 2001; Arnoldi et al. 2018; Medeiros et al. 2025) and experimentally (Amor et al. 2020) to the responses of natural systems to pulse disturbances, and some to press disturbances (Morozov et al. 2020; Inamine et al. 2022). But previous work has not noted or exploited a direct connection between pulses and a press. In discrete-time models, a press disturbance can be viewed simply as a continuing series of pulses, one in every time interval. In continuous-time models a pulse is a short-lived disturbance, whereas a press is just a long-lived ongoing version of that pulse. Here, we use that equivalence to link transient response after a pulse to the response to a press.

To set the stage, we provide a brief overview of structured populations in discrete time, represented by a matrix population model (MPM) (Caswell 2001). The MPM, denoted by **B** has elements *b*_*ij*_ that describe the per-capita contribution from state *j* to state *i* in one time step. These transition rates are non-negative and may combine multiple vital rates (*e*.*g*., fertility is often the product of survival and fecundity). The long-run growth rate of the population is given by the dominant eigenvalue of **B** and is denoted by *λ*_0_. There are two eigenvectors associated with *λ*_0_. The right eigenvector **u**_0_ (i.e., **Bu**_0_ = *λ*_0_**u**_0_) yields the stable stage distribution (SSD), that is, the proportion of individuals across states in the long run. The left eigenvector **v**_0_ gives the stable reproductive value, which describes the contribution of individuals to future reproduction. Together, (*λ*_0_, **u**_0_, **v**_0_) describe the asymptotic behaviour of the population.

Now that we have described the long-run behaviour, we ask what happens when a small disturbance (such as a fire) shifts the system away from equilibrium. To examine this, we assume that the population is at its SSD and subject it to a one-time pulse disturbance. We define a disturbance as changing the population’s transition rates between states (ages or stages), and not as directly affecting population structure. Thus, at time *t* = 0, the population is at **u**_0_ and is impacted by a pulse between time *t* = 0 and *t* = 1. The result is that the subsequent population structure, at time *t* = 1, is no longer the SSD, but is a sum, 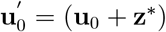, where the pulse produces the effect **z**^*∗*^ (see Methods section). The pulse does not recur (by definition), so its effect diminishes over time. So, the population structure at time *t* = 2 is the sum of the SSD and the 1-time diminished effect of the original pulse. Similarly, at time *t* = 3, the population structure is the sum of the SSD and the 2-time diminished effect of the original pulse. Eventually, the effect decays and the structure converges to the original stable structure **u**_0_.

In contrast, the response to a press (i.e., ongoing) disturbance (such as an ongoing drought) is an accumulation of responses to repeated pulses. The initial state is the SSD at time *t* = 0, and a pulse between time 0 and 1 produces at time 1 the change **z**^*∗*^, exactly as before (as shown in Figure 1). At time *t* = 2, however, we add together two effects: the time diminished effect of the first pulse (between time 0 and 1) and the next pulse (between time 1 and 2). At time *t* = 3, we again add the new pulse to the remaining effect of previous pulses; the sum is the effect of the press disturbance. In the long-term, we have a cumulative sum that yields the deviation from the original SSD to a new SSD; the cumulative sum is given by equation 9, and illustrated in general by Figure 1. Numerical examples are in Figures 2a and 2b.

**Figure 1:**
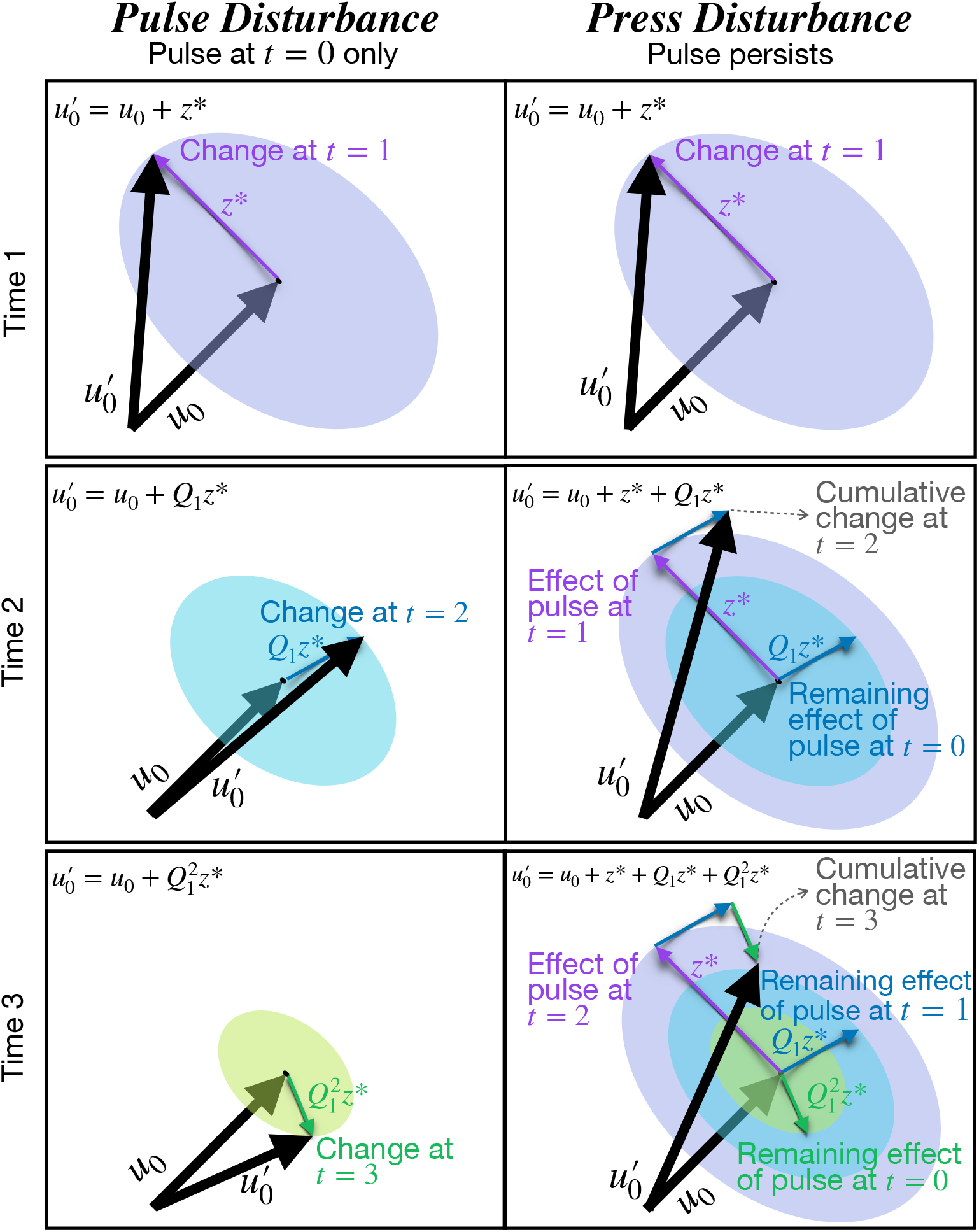
Dynamics of population structure after a pulse and press. The left panels illustrate transient dynamics of a structured population after a pulse (one-time at *t* = 0) disturbance. At time *t* = 1, the purple arrow depicts a pulse disturbance acting on the stable distribution **u**_0_, producing a deviation **z**^*∗*^. Here, **z**^*∗*^ denotes the perpendicular shift in the population structure following the disturbance. At time *t* = 2, the pulse decays showing subsequent rotation and shrinkage of deviations from the SSD **u**_0_ (blue arrow), and then at time *t* = 3 (green arrow), *etc*.. The disturbance decays according to 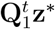, where **Q**_1_ matrix governs the rate of decay and the transient dynamics. The blue vector represents the 1-time diminished disturbance, and the green vector represents the 2-time diminished disturbance. Eventually, the structure converges to original stable structure **u**_0_ in the bottom left panel. In the right panel, results of a press (permanent) disturbance are illustrated as the sum of the effects of the current pulse (purple vector), and the decaying effects of past pulses (blue and green vector at time *t*− 1 and *t*− 2, respectively). In the case of press disturbance (right panel), the convergence is to a new stable structure 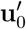.

**Figure 2a:**
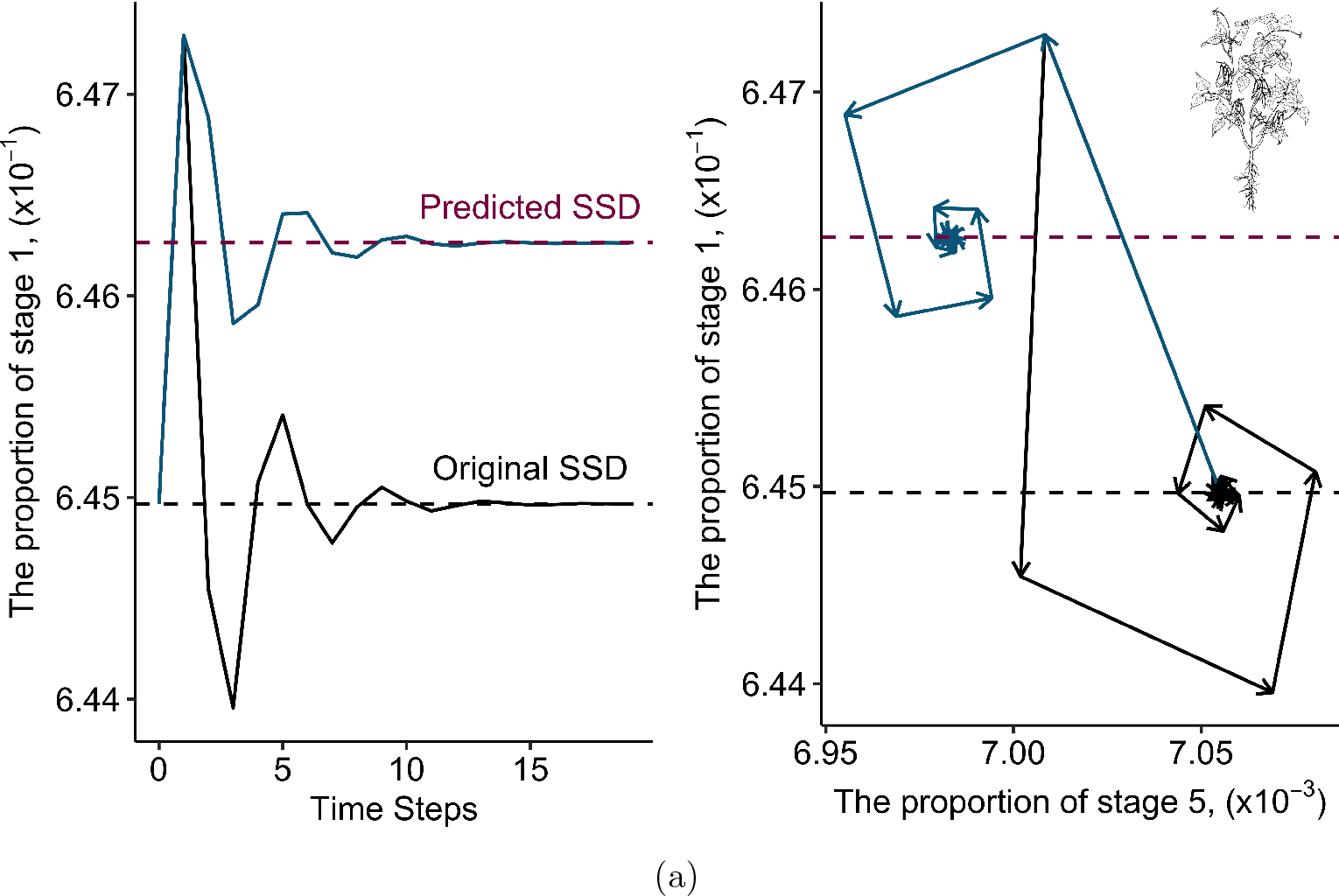
Comparing the response to pulse and press for *Phaseolus lunatus*. The left panel shows transient dynamics in Stage 1 after a pulse (black line) or a press (blue line) disturbance to the (1,6) and (6,5) elements of the matrix population model. Stage 1 converges back to the original stable stage distribution (SSD; dashed black line) following a pulse disturbance, while a press disturbance leads to convergence toward a new SSD (dashed magenta line), analytically predicted using the Transient Response Matrix (TRM). On the right panel, the transient dynamics are visualized in a phase plane with Stage 5 on the x-axis and Stage 1 on the y-axis. Arrowheads indicate the direction of convergence. Both stages converge to the original SSD (black triangle) after a pulse disturbance, while a press disturbance drives convergence to a distinct SSD. Lima Bean sketch credit: IBPGR (1982).

**Figure 2b:**
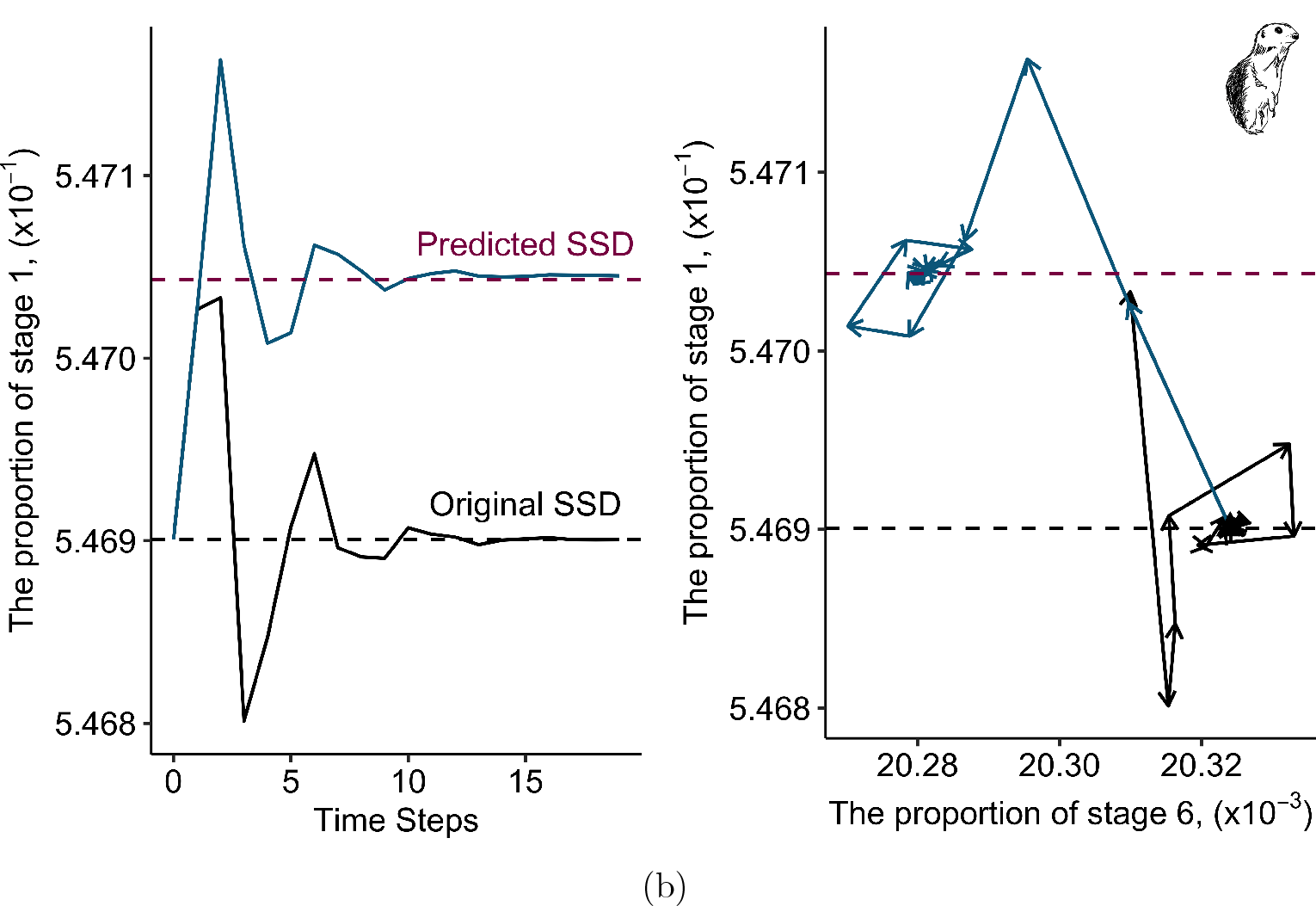
Comparing the response to pulse and press for *Spermophilus dauricus*. The left panel shows transient dynamics in Stage 1 after a pulse (black line) or a press (blue line) disturbance to the (1,7) and (7,6) elements of the matrix population model. Stage 1 converges back to the original stable stage distribution (SSD; dashed black line) following a pulse disturbance, while a press disturbance leads to convergence toward a new SSD (dashed magenta line), analytically predicted using the Transient Response Matrix (TRM). On the right panel, the transient dynamics are visualized in a phase plane with Stage 5 on the x-axis and Stage 1 on the y-axis. Arrowheads indicate the direction of convergence. Both stages converge to the original SSD (black triangle) after a pulse disturbance, while a press disturbance drives convergence to a distinct SSD. Daurian squirrel sketch credit: Harman Jaggi.

Our framework shows how transient responses to pulse disturbances lead to a quantitative description of press disturbances. We derive the connection from pulses to presses via what we label a Transient Response Matrix (TRM). We apply our framework to two stage-structured matrix population models (*Phaseolus lunatus* and *Spermophilus dauricus*) from the COMADRE and COMPADRE database (Salguero-Gómez et al. 2015, 2016). Using these examples, we show that small perturbations (pulse and press) leave distinct and predictable signatures on the population structure. The response to a small pulse and press perturbation matches theoretical predictions exactly (Figure 2a and 2b). Thus, our framework provides a way to link experimental or observational manipulations directly to fitness consequences. For example, a researcher testing how invasive plants alter resource availability (press) or how an extreme frost event reduces survival in a single year (pulse) could use our approach to quantify both within the same modelling structure. The examples highlight the utility of our analysis in linking transient dynamics with long-term demographic consequences across different life-histories.

Interestingly, the TRM also yields the second derivatives of the population growth rate with respect to demographic rates (matrix elements). Caswell (1996) and Shyu and Caswell (2014) have long argued for the importance of second derivatives and developed methods for their computation. Whilst valuable, these methods have heavily relied on vector calculus-potentially obscuring important demographic processes connected to second-order derivatives. Here we offer an intuitive method using perturbation theory and our approach is valid for any discrete-time, age or stage-based population model.

Previous work (Jiang et al. 2022) has identified a positive association between life-history metrics and time to convergence. Based on previous macroecological work, we hypothesized a correlation between TRM and generation time (average age of survival-weighted reproduction). Using 439 unique species, we indeed find that a positive association between dominant (nonzero) eigenvalue of TRM and the generation time (in the Results section). In addition, the TRM is directly related to the cumulative distance to stability as defined by Cohen (1977). In the next section, we define matrix population models (MPMs) and summarize (known) effects of pulse disturbances, and our approach to transient dynamics. Then we show precisely how pulses add to a press and how the responses to a press depend on the transient response matrix (TRM), **J**_0_.

## Methods

Table 1 provides a summary of key definitions used for MPMs and its decomposition (as we discuss later in this section). As outlined in the Introduction, we consider structured population models in discrete-time described by a MPM. The matrix may be a discretized version of an integral population model. The unperturbed MPM is denoted by **B**, and its elements (*b*_*ij*_) are transition rates (per-capita contributions of individuals) from state *j* at time *t* to state *i* at time (*t* + 1). The matrix **B** is non-negative since transition rates ≥ 0. Starting with an initial population vector **n**(0) and following the population at time *t*, **n**(*t*), yields **n**(*t*) = **B**^*t*^**n**(0), over time *t* (Caswell 2001).

**Table 1:**
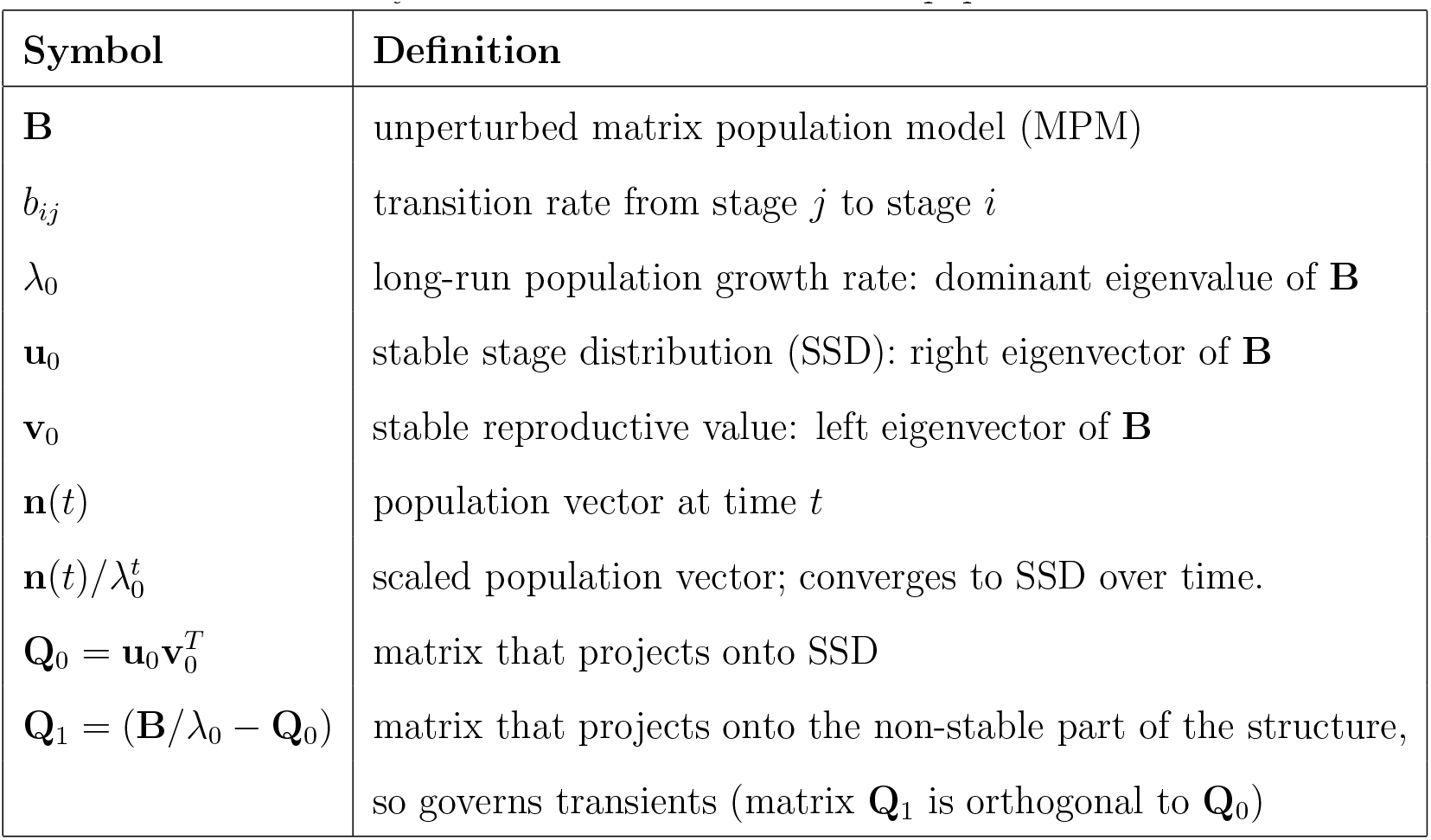
Symbols and definitions for matrix population models.

| Symbol | Definition |
| --- | --- |
| $\mathbf{B}$ | unperturbed matrix population model (MPM) |
| $b_{ij}$ | transition rate from stage $j$ to stage $i$ |
| $\lambda_0$ | long-run population growth rate: dominant eigenvalue of $\mathbf{B}$ |
| $\mathbf{u}_0$ | stable stage distribution (SSD): right eigenvector of $\mathbf{B}$ |
| $\mathbf{v}_0$ | stable reproductive value: left eigenvector of $\mathbf{B}$ |
| $\mathbf{n}(t)$ | population vector at time $t$ |
| $\mathbf{n}(t)/\lambda_0^t$ | scaled population vector; converges to SSD over time. |
| $\mathbf{Q}_0 = \mathbf{u}_0 \mathbf{v}_0^T$ | matrix that projects onto SSD |
| $\mathbf{Q}_1 = (\mathbf{B}/\lambda_0 - \mathbf{Q}_0)$ | matrix that projects onto the non-stable part of the structure,<br>so governs transients (matrix $\mathbf{Q}_1$ is orthogonal to $\mathbf{Q}_0$) |

When the population projection matrix **B** is primitive and irreducible (as is typical for ecological models such as Leslie or Lefkovitch matrices), there is a dominant eigenvalue *λ*_0_ with associated right and left eigenvectors, **u**_0_ and **v**_0_, respectively (Caswell 2001). The dominant eigenvalue *λ*_0_ determines the asymptotic population growth rate. The right eigenvector **u**_0_ represents the stable stage distribution (SSD) toward which the population converges in the long term, and the left eigenvector **v**_0_ defines the stable reproductive value describing their relative contribution to future population growth. The SSD **u**_0_ is scaled so that its entries sum to one, expressing the proportion of individuals in each stage. This gives **e**^*T*^ **u**_0_ = 1, where **e** is a vector of ones. The reproductive value vector **v**_0_ is scaled so that the total reproductive value of the SSD equals one: 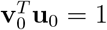. Here, *T* is transpose of a matrix.

We use two published MPMs from publicly available databases COMPADRE and COMADRE (Salguero-Gómez et al. 2015, 2016). The first MPM is a 6-stage annual plant for *Phaseolus lunatus* (Lima bean), with a population growth rate *λ*_0_ = 0.76. The second is a 7-stage mammalian matrix for *Spermophilus dauricus* (Daurian ground squirrel) with a population growth rate *λ*_0_ = 0.92. In the results seciotn, we will use these contrasting life-histories (an annual plant and a mammal) to illustrate our framework and demonstrate the generality of our results.

### Decomposing population projection matrix into stable and transient components

To analyze transient and asymptotic behaviors, we rescale the original matrix by dividing by the population growth rate *λ*_0_. Next, we decompose the rescaled matrix 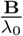 into two parts:

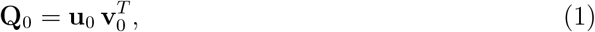

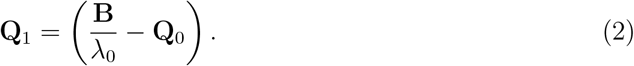

The matrix **Q**_0_ comprises of stable vectors (**u**_0_ and **v**_0_) and projects any population vector onto the SSD, capturing the long-term stable dynamics. Because 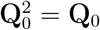, it is a projection matrix. Moreover, the matrix **Q**_1_ is orthogonal to **Q**_0_ since **Q**_0_**Q**_1_ = **Q**_1_**Q**_0_ = 0.

This leads to the full decomposition:

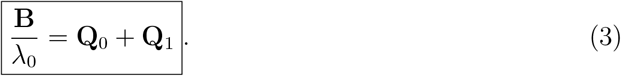

Using this decomposition in equation (3) and the fact that **Q**_0_ and **Q**_1_ are orthogonal, the population structure at time *t*, scaled by 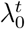 is given by:

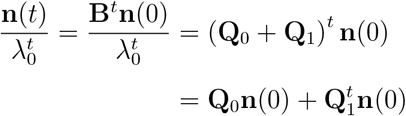

The term **Q**_0_**n**(0) = (**v**_0_, **n**(0)) **u**_0_ gives the asymptotic SSD since it is a scalar multiple of **u**_0_. The term 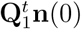 governs the transient trajectory. This is because the spectral radius of **Q**_1_ is less than one, and thus the powers of **Q**_1_ vanish over time, implying convergence to the SSD:

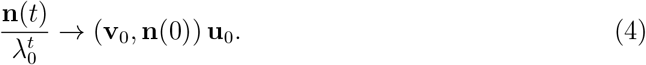

The decomposition in equation (3) separates the stable (**Q**_0_) and unstable (**Q**_1_) components of population projection matrix **B**. This framework underpins recent ecological applications studying transient dynamics (Koons et al. 2005; Haridas and Tuljapurkar 2007; Capdevila et al. 2021; Jiang et al. 2022).

### Transient dynamics after a pulse disturbance

Note that Table 2 summarizes the key definitions related to transient dynamics and perturbation analysis. Let us assume a population is initially at its SSD, **u**_0_, with associated stable reproductive value (SRV) vector **v**_0_, and projection matrix **B**. If a pulse perturbation acts at *t* = 0 for one time step, the population is no longer at equilibrium. To characterize the effects of this deviation, we use a decomposition based on the eigenstructure of **B**.

**Table 2:**
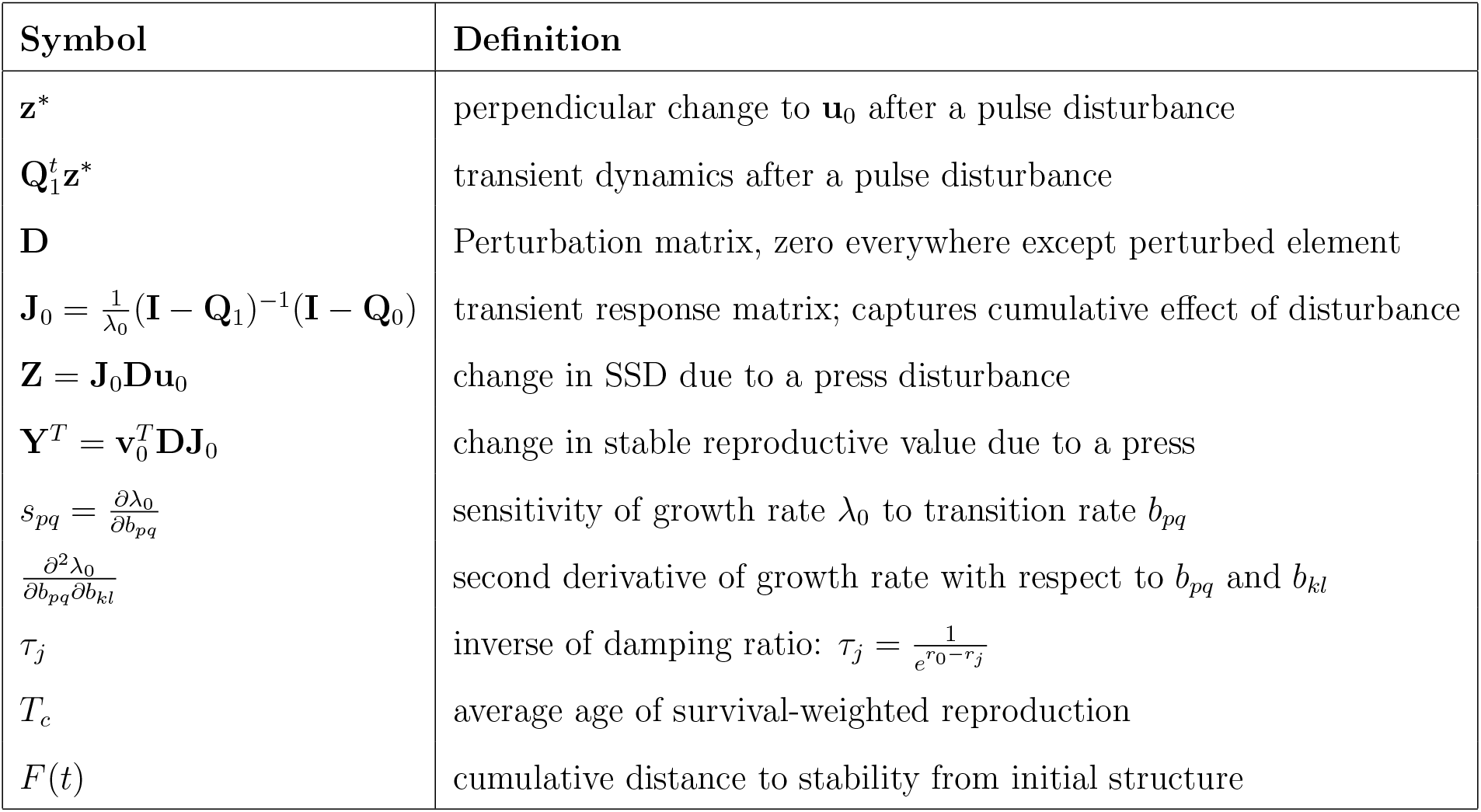
Symbols and definitions for perturbation analysis.

A small pulse perturbation modifies the matrix **B** to **B** + *ϵ***D** for one time step, where *ϵ* is the magnitude of pulse perturbation. The matrix **D** represents the matrix element being perturbed and is zero everywhere except at the entry (**D**_*ij*_) being perturbed. In ecological terms, this corresponds to a temporary change in a demographic rate. For example, perturbing the fertility rate *b*_15_ would make the element **D**_15_ = 1 as the only nonzero element of **D**. At time *t* = 1, the population structure is no longer **u**_0_, but instead becomes

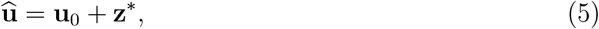

where the shift **z**^*∗*^ lies in the transient subspace orthogonal to the SSD:

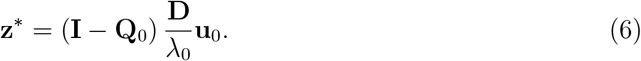

Because **z**^*∗*^ lies in the transient subspace, it determines how the population structure shifts from and eventually returns to the SSD. The proof for equation (6) is given in Appendix A.2.

After the pulse ends, the population is again governed by the original matrix **B**, and hence evolves according to:

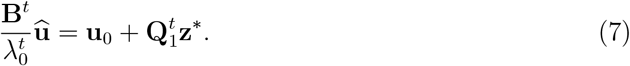

As discussed previously:

- The term **u**_0_ remains constant-the population converges back to the **u**_0_ because powers of **Q**_1_ decay to zero.
- The term 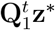 evolves over time and captures the transient dynamics.

The path taken during the convergence to **u**_0_ is determined by the transient component **z**^*∗*^ and its successive projections 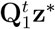. The full trajectory of the population structure through time (after a pulse) is the sequence:

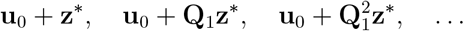

Each step in this sequence represents the population structure at time *t* = 1, 2, 3, … after the pulse. The rate and direction of convergence depend on the structure of **Q**_1_ and the form of **z**^*∗*^. The term 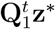 both shrinks in magnitude and may rotate in direction (as some directions may decay faster than others), depending on the eigenstructure of **Q**_1_. Thus, **Q**_1_ is the time-dimished effect of the pulse produced by disturbance **z**^*∗*^ as discussed in the Introduction. We illustrate the key finding from this subsection in the left panel of Figure 1.

Our approach separates stable and transient components, offering a biologically interpretable approach to pulse dynamics. The research builds on and extends previous work on transient dynamics (Haridas and Tuljapurkar 2007; Neubert et al. 2002; Koons et al. 2017; Stott 2016).

### Press disturbance as a series of pulses

A press disturbance refers to a permanent change to the population projection matrix **B**, represented as **B** + *ϵ***D**, where *ϵ* is the magnitude of perturbation and **D** is a matrix with nonzero entries for elements that perturbed. We assume the perturbation is small, so that first and second order approximations apply. A key idea is that a press disturbance is equivalent to an ongoing sequence of identical pulse perturbations.

We begin by characterizing the response to a single pulse, which shifts the structure by a vector **z**^*∗*^ perpendicular to **u**_0_, as described in equation (6) given by 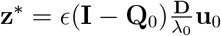. This vector satisfies **Q**_0_**z**^*∗*^ = 0 and captures the deviation from the SSD caused by the disturbance.

Because a press perturbation acts at each time step, the full trajectory of the population structure consists of accumulating transient effects and new perturbations. At time *t* = 1, the structure is **u**_0_ + **z**^*∗*^ (same as pulse disturbance). Now at *t* = 2, we apply the new perturbation and add the decayed contribution of the previous perturbation:

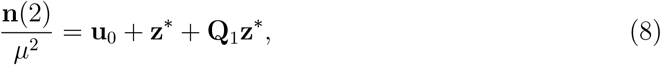

where *µ* is the new dominant eigenvalue under the press disturbance. That is, *µ* is the dominant eigenvalue of the **B** + *ϵ***D**. Continuing in this way, we apply the new perturbation and add the decayed contribution of the previous perturbations to get the general expression:

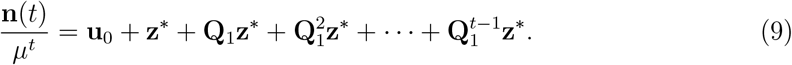

We illustrate this approach in the right panel of Figure 1. As time increases (*t* → ∞), the cumulative effect converges to a geometric series:

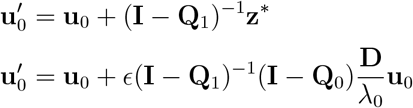

where 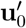 represents the new stable stage distribution (SSD) under the press disturbance. Similarly (as discussed in Appendix B), an analogous argument can be made for the stable reproductive value.

Next, we define the Transient Response Matrix (TRM) as follows:

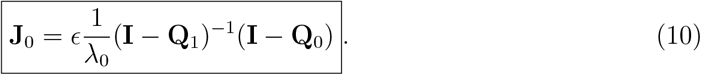

Then the change in SSD (upto second order) from the press disturbance becomes:

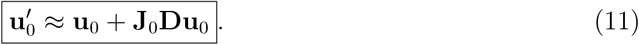

These results provide an approximation for the effects of press disturbances on the long-term population structure and reproductive values. The approximation for 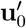 and 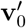 is exact in the limit of small perturbations since higher-order terms can be neglected. This means that we have obtained the linear change (equivalently, the first derivative) of the SSD (or SRV).

Our approach uses only the dominant eigenstructure and the TRM **J**_0_ thereby generalizing the analysis in Caswell (1996) and making it broadly applicable to structured population models. The transient response matrix **J**_0_ maps small changes in the projection matrix to changes in the SSD (or SRV). The TRM **J**_0_ describes the accumulation of the effects of continued disturbance, as well as the intrinsic ability to dampen the perturbations. In a later section, we discuss the many applications of the new matrix **J**_0_.

We illustrate the dynamics of pulse (left panel) and press disturbances (right panel) in Figure 1, which shows how transient deviations decay and evolve over time using colored time-diminished vectors. Further, we test our theory on two life-histories as shown in Figure 2a and Figure 2b in the Results section.

### Quantifying change in fitness: sensitivity and nonlinear response

In MPMs, fitness is the long run growth rate *λ*_0_ determined by the elements of matrix **B**. Ecologists have long studied the sensitivities *s*_*pq*_ of fitness (Caswell 2001) with respect to each matrix element *b*_*pq*_. We begin with a simple intuitive approach to sensitivity, and then use it to study the second derivative (or curvature) of fitness (Kajin et al. 2025).

#### Linear change and sensitivity

Consider a stable population at the SSD, with a fraction *u*_0*q*_ of individuals in stage *q*, and a one-period population growth rate of *λ*_0_. Hence *u*_0*q*_ is the fraction of individuals available to make a *q* → *p* transition. Since *b*_*pq*_ is the rate for that transition, the number of individuals in state *p* produced by that transition is proportional to the product *u*_0*q*_ *b*_*pq*_. In that final state, every individual has reproductive value *v*_0*p*_, so the product

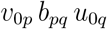

is the relative contribution of the *q* → *p* transition to population growth. The total of these contributions over all initial and final states is *λ*_0_ (as it should be).

Now suppose that we add a small amount *∂b*_*pq*_ to the rate for the *q* → *p* transition, i.e., we change (*b*_*pq*_) to (*b*_*pq*_ + *∂b*_*pq*_). Following the logic above, the change in growth rate is the product of three terms:

a. the fraction of population that is subject to this state change, i.e., *u*_0*q*_;
b. the change in that transition rate, i.e., *∂b*_*pq*_;
c. the relative “value” of an additional contribution to final state *q*, i.e., *v*_0*p*_.

The product is *∂b*_*pq*_ *v*_0*p*_ *u*_0*q*_. Dividing by *∂b*_*pq*_, we conclude that the first derivative of population growth rate is

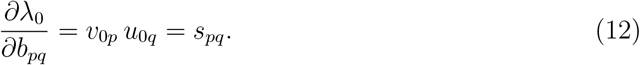

Here, *s*_*pq*_ is the sensitivity with respect to *b*_*pq*_ element and the last equality is the standard result (Caswell 2001). We now use our new simple method to find the second derivatives of fitness.

#### Second-order fitness response to matrix perturbation

Suppose a press disturbance affects the *b*_*pq*_ and *b*_*kl*_ elements of the matrix **B**, where *p, q, k, l* are indices. For example, a disturbance to the fertility element *b*_15_ would represent a shift in offspring production, whereas a disturbance in survival element *b*_21_ would correspond a change in state survival. Figure 3 illustrates two sequences of applying these changes. In the first path, we modify the (*kl*) transition rate by *f* (I → II), followed by the (*pq*) rate by *g* (II → IV). In the second path, we reverse the order: modify (*pq*) first (I → III), then (*kl*) (III → IV). Both routes must yield the same total change in fitness, as fitness is a scalar function of matrix entries.

**Figure 3:**
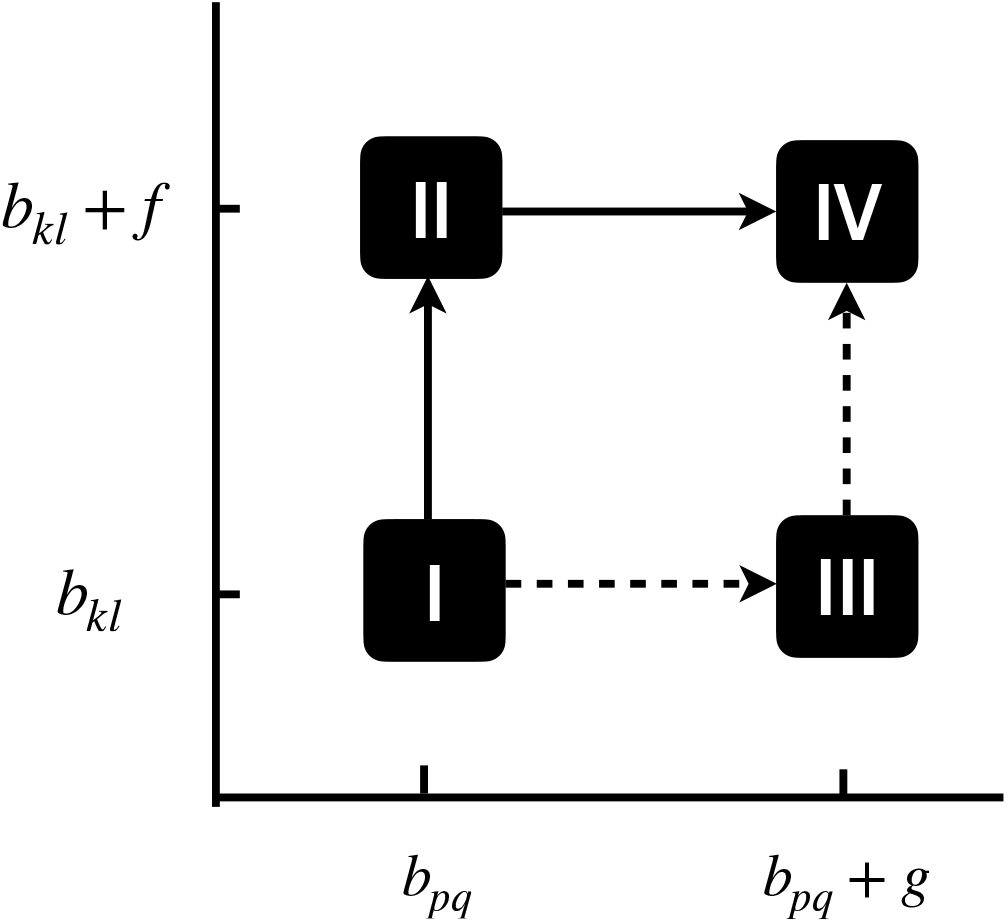
Computing the second derivatives of fitness with respect to matrix elements. The horizontal and vertical axes indicate rates for the two demographic transitions, (*p, q*) and (*k, l*). Point I indicates the starting values, where the fitness is *λ*_0_, SSD is **u**_0_ and stable reproductive value is **v**_0_. A press disturbance of both demographic rates ends at point IV. We consider two possible routes. First route: go from I to II by changing only the demographic rate for the *k* ← *l* transition (*i*.*e*., *b*_*pq*_ is unchanged but *b*_*kl*_ becomes *b*_*kl*_ + *f*). At II the new stable population is, say, **u**_0_ + **Z**_1_ and the new reproductive value is, say, **v**_0_ + **Y**_1_. Next go from II to IV, by changing only *b*_*pq*_ by an amount *g*. Second route: starting at I, go from I to III by perturbing the *p* ← *q* transition rate by an amount *g*. At III the new stable population is, say, **u**_0_ + **Z**_2_ and the new reproductive value is, say, **v**_0_ + **Y**_2_. Next, go from III to IV by changing only *b*_*kl*_ by an amount *f*.

We first consider the I → II → IV path. At point I, the SSD is **u**_0_ and the SRV is **v**_0_. Using the logic from the previous subsection, the fitness change in the I → II step is:

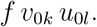

At point II, the SSD has changed from **u**_0_ to **u**_0_ + **Z**_1_, where **Z**_1_ quantifies the first-order change in SSD due to the perturbation in *b*_*kl*_ (as discussed in Appendix B):

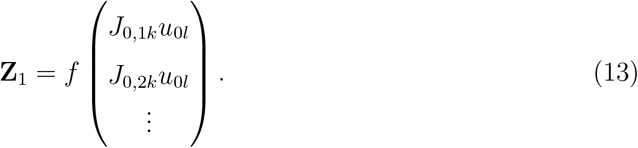

Also at II, the stable reproductive value is (**v**_0_ + **Y**_1_) with

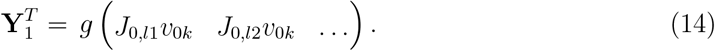

So in the transition II to IV, the fitness changes by the product of

a. the stable proportion in stage *q*, which equation (13) shows is (*u*_0*q*_ + *Z*_1*q*_),
b. the change in the rate, *g*,
c. the stable reproductive value in stage *p*, which equation (14) shows is (*v*_0*p*_ + *Y*_1*p*_).

Thus the total change in the presses from I to II, and II to IV (see Figure 3) is

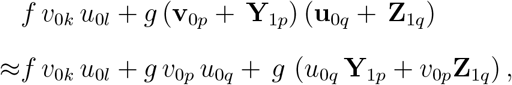

where the higher order term *g***Y**_1*p*_**Z**_1*q*_ is ignored because it is close to zero. Split this up into two bits,

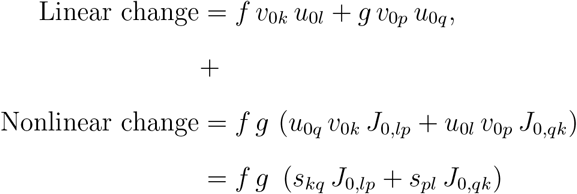

When *f* and *g* are small, the nonlinear change yields the second derivatives of fitness:

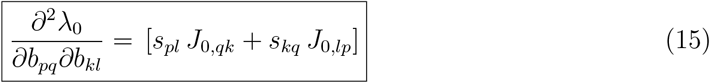

where we use the sensitivities *s*_*kq*_ = *v*_0*k*_ *u*_0*q*_ and *s*_*pl*_ = *v*_0*p*_ *u*_0*l*_. This expression for the second derivative is symmetric with respect to an exchange of the elements *b*_*pq*_, *b*_*kl*_ (as it should be). The curvature of fitness is measured by the second derivatives in equation (15) and depends on both the first-order sensitivities and the TRM (**J**_0_). As shown in Fig 3, we could alternatively go from I to III and then III to IV. That process involves different changes to the SSD and reproductive value, but yields the same final result. The details of this calculation are presented in the Appendix B.3. In Appendix B.4, we provide a stepwise calculation of second derivatives for *Phaseolus lunatus*. Our calculation for the final answer exactly matches Shyu and Caswell (2014), however our approach is more intuitive and based on application of perturbation theory. A complete derivation can be found in Appendix B.2.

### Transient response matrix J_0_

The previous two sections identify the central role of the TRM **J**_0_ in the linear and nonlinear responses to a disturbance, where,

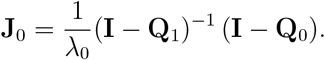

Since matrix **Q**_1_ shapes the transient dynamics after a pulse, those transient dynamics also shape the TRM, and thus the nonlinear change in growth rate. This connection between transients and the nonlinearity of the dominant eigenvalue is unexpected and intimate. Here, we further illuminate this connection by three new results that connect the TRM to transients.

### TRM and cumulative distance to stability

In general, even when the MPM does not have distinct eignevalues, an important metric to assess the difference between an observed stage distribution and the SSD is the cumulative distance to stability (Cohen 1979a). Several studies (Williams et al. 2011; White et al. 2013) have employed this metric. Here we show that the TRM is closely related to the cumulative distance.

Any non-stable initial population distribution at time *t* = 0 converges towards the SSD. Following Cohen (1979a), at each time *t >* 0, the “distance” from stability is measured by summing the elements of the difference vector 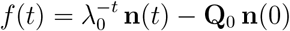. Convergence means that this distance is decreasing, so the cumulative vector 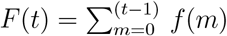 should have a limit. For an initial population vector **n**(0) Cohen showed this limit to be

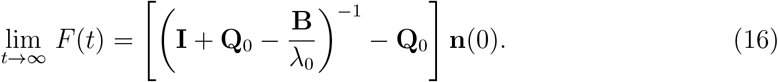

But we find (see Appendix D for matrix equivalence) that Cohen’s cumulative distance is just

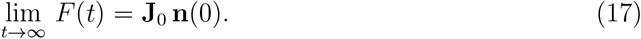

Thus the TRM **J**_0_ is directly related to the asymptotic cumulative distance to stability.

### TRM for population matrices with distinct eigenvalues

Suppose that the MPM **B** has distinct eigenvalues, so that **u**_*j*_, **v**_*j*_ are the right, left eigenvectors for eigenvalue 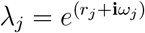 for *j* ≥ 1. Below, superscript *†* indicates a complex conjugate transpose.

Define the eigenvalue ratios

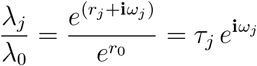

where 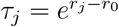 and *τ*_*j*_ *<* 1; the *τ*_*j*_ are the inverse damping ratios Caswell (2001). Then a spectral decomposition and the definitions (1) and (2) yield the expression:

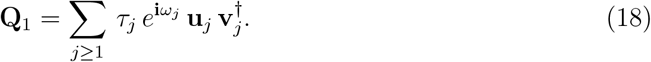

From this, we can now expand **J**_0_ (see Appendix C) as

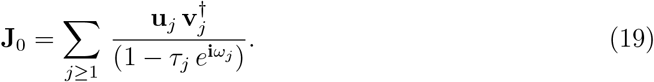

This is a spectral decomposition of the TRM (excluding the eigenvector **u**_0_ for which the eigenvalue is 0). Thus the vectors **u**_*j*_, **v**_*j*_ are right, left eigenvectors of **J**_0_ corresponding to eigenvalue 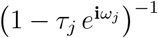. One consequence is that the dominant non-zero eigenvalue of the TRM depends on the damping ratio *τ*_1_, being low when *τ*_1_ ≪ 1 and becoming large when *τ*_1_ approaches 1.

## Results

To test our framework and evaluate how structured populations respond to disturbances, we analyze both short-term (pulse) and long-term (press) perturbations for two population projection matrices. Our first result shows that pulses and presses leave distinct signatures on population structures and can be predicted using our method. Next, we examine how the transient response matrix (TRM, **J**_0_) relates to life-history traits like generation time (*T*_*c*_), defined as average age of survival-weighted reproduction. Finally, we demonstrate how first- and second-order derivatives of population growth rate vary in direction and magnitude. Together, our results show how transient dynamics are linked to long-term outcomes and life-history traits via transient repsonse matrix. Note that a table of key vectors and matrices with their definitions is provided in Tables 1 and 2.

### Convergence to stable stage under press and pulse disturbances for two life-histories: *Phaseolus lunatus* and *Spermophilus dauricus*

To examine how press and pulse disturbances shape population structure over time, we simulate perturbations for two life histories: *Phaseolus lunatus* (Lima bean), an annual plant with a 6-stage MPM, and *Spermophilus dauricus* (Daurian ground squirrel), a mammal represented by a 7-stage MPM. The MPMs taken from open source COMPADRE and COMADRE databases, respectively (Salguero-Gómez et al. 2015, 2016). We focus on disturbances applied to fecundity and survival rates, specifically elements (1,6) and (6,5) for Lima bean (Figure 2a); (1,7) and (7,6) for Daurian ground sqquirrel (Figure 2b). The perturbations reflect environmental shocks that have one-time (pulse) or persistent effect (press) on the two life-histories.

In the left panel of Figure 2a and 2b, we track the proportion of individuals in Stage 1 over time following either a single-time pulse (black line) or a sustained press (blue line) disturbance (increased each by 2%). The system returns to the unperturbed stable stage distribution (SSD) after a pulse, as expected from matrix analysis discussed above. In contrast, the press disturbance leads to a new equilibrium structure, indicated by the magenta dashed line. The new SSD is accurately predicted by the TRM **J**_0_ = (**I** − **Q**_1_)^−1^(**I** − **Q**_0_) using perturbation theory, reinforcing its utility in forecasting stable st(ages) under sustained environmental pressures.

The right panel projects the same dynamics onto a phase plane composed of Stage 1 and Stage 5 proportions for Lima bean; Stage 1 and Stage 6 for Daurian ground squirrel. Both trajectories exhibit damped convergence—either toward the original SSD (black triangle) for the pulse or a new SSD under press, confirming theoretical expectation on how the long-run structure shifts under press perturbations. The convergence trajectories spiral back to stable state, revealing differences in the strength and shape of transient dynamics across the two species.

We numerically verify that even moderate perturbations to MPMs yield excellent agreement between the predicted and observed 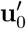, demonstrating the utility of the approximation as shown in Figure 2a and Figure 2b. Our result illustrates how small, short-lived disturbances can generate complex trajectories in stage structure before long-term convergence as shown in Figure 2a and Figure 2b.

### Linking TRM to life-history traits

Jiang et al. (2022) found a strong correlation between damping time and generation time (*T*_*c*_), a key life history trait (Gaillard et al. 2005). That finding led us to hypothesize that the dominant non-zero eigenvalue of TRM (**J**_0_) is also correlated with generation time (*T*_*c*_). To examine this hypothesis, we analyzed 439 unique age and stage-structured species (after correcting for phylogenetic inertia) using the COMADRE Animal Matrix Database (Salguero-Gómez et al. 2016), the COMPADRE Plant Matrix Database (Salguero-Gómez et al. 2015), and previously published mammalian database in Jiang et al. (2022). As shown in Figure 4, we find that the dominant non-zero eigenvalue of **J**_0_ is indeed strongly correlated with generation time *T*_*c*_ on the log-log scale.

**Figure 4:**
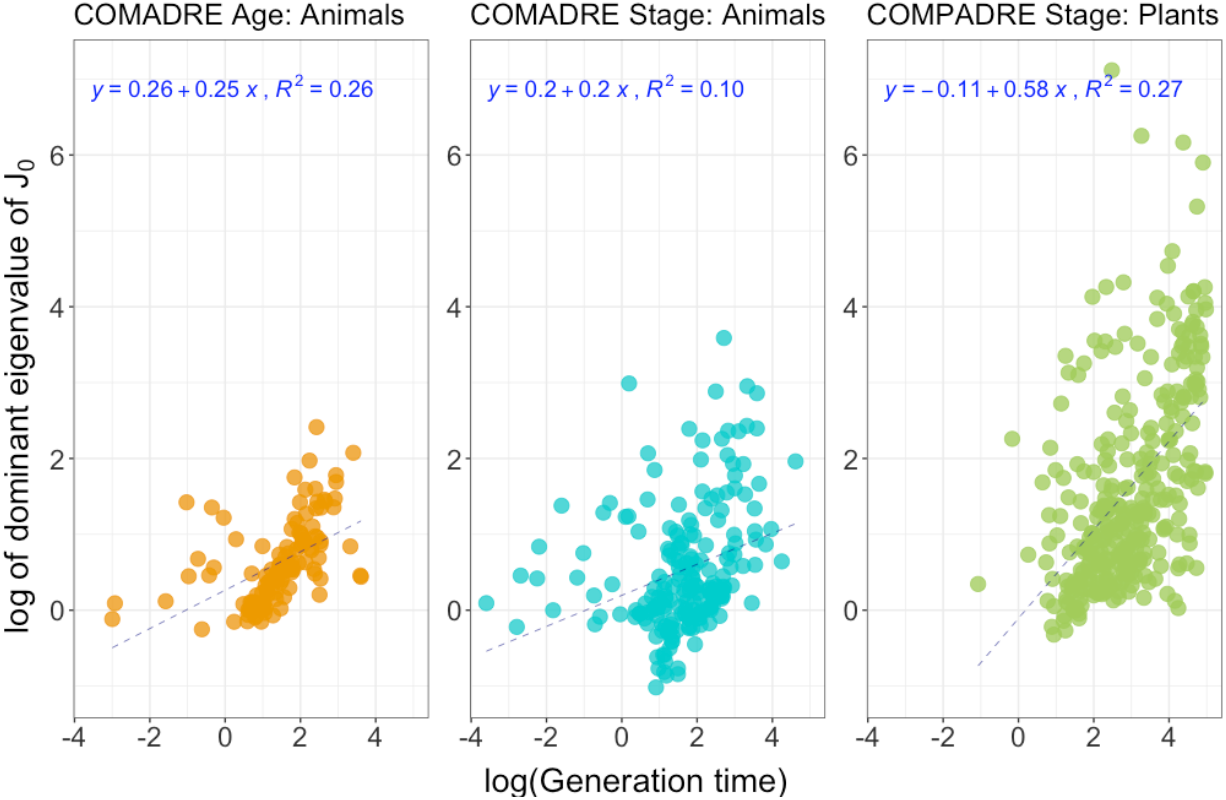
Relationship between generation time and TRM. The figure examines the relationship between generation time (*T*_*c*_) and the dominant eigenvalue of **J**_0_ based on phylogenetic generalized least squares. The y-axis corresponds to log of the dominant eigenvalue of **J**_0_ and the x-axis corresponds to log of generation time *T*_*c*_. Each panel corresponds to a database: COMADRE age-structured matrices, COMADRE stage-structured matrices, COMPADRE stage-structured matrices. The correlation between dominant eigenvalue of **J**_0_ and *T*_*c*_ is consistently positive across all databases.

This relationship can be understood as follows: species with longer generation times exhibit slower convergence to their stable distribution following a perturbation. Since the TRM **J**_0_ captures the cumulative effect of transient dynamics under disturbance, its dominant eigenvalue effectively summarizes the timescale of decay toward equilibrium. A large generation time *T*_*c*_ implies that demographic transitions are inherently slower, which delays convergence and amplifies the transient contribution.

### Comparing range of sensitivities, TRM and second-order derivatives of population growth rate

We examine the range and distribution of key quantities: the second derivatives of population growth rate, the Transient Response Matrix (TRM, **J**_0_) and first-order sensitivities for *Phaseolus lunatus*. As shown in Figure 5, both **J**_0_ and the second derivatives exhibit a wide spread of values, spanning positive and negative ranges. Ecologically, this means that small perturbations to survival or fertility can either increase or decrease fitness depending on the location and direction of perturbation, and how it interacts with the structure of the matrix model. Therefore, positive values can amplify transient deviations, whereas negative values can dampen deviations from the stable state.

**Figure 5:**
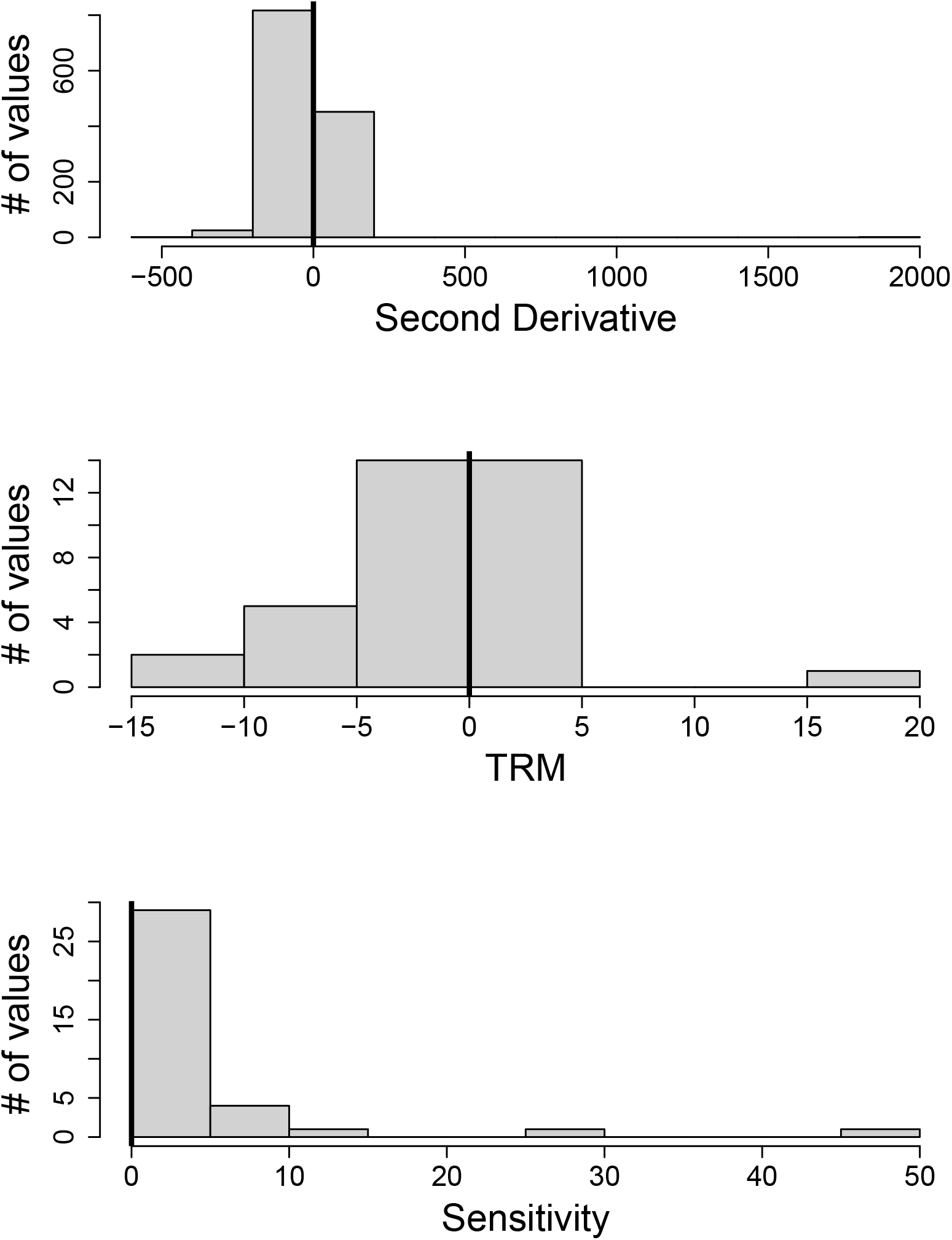
Histograms to show range and distribution in values of second deriva-tives, TRM (J_0_), and sensitivities. Second derivative of population growth rate and **J**_0_ both have positive and negative values, but sensitivity of growth rate only have positive values.

In contrast, the first-order sensitivities are strictly non-negative, because all terms in *s*_*pq*_ = *v*_0*p*_*u*_0*q*_ are always positive for population projection matrices. The new insight from second derivatives and TRM is that they capture the nonlinear and potentially deteriorating effect of perturbations. This is because the curvature, captured by the second derivative of population growth rate can be positive or negative based on effect of perturbation. Thus, second derivatives and TRM extend classical sensitivity analysis, emphasizing the importance of second-order terms in forecasting demographic responses to disturbance.

## Discussion

Population ecology has made important progress in understanding how natural populations respond to one-off (*i*.*e*., pulse) disturbances (Jentsch and White 2019; Tao et al. 2021). Here, we developed a novel approach to examine how natural populations respond to press disturbances, which can be expressed as repeated pulses, when the disturbances are small. Our results reveal an intimate and unexpected connection between the nonlinearity of population growth rate to vital rates, and transient dynamics. We apply our theoretical predictions to *Phaseolus lunatus* and *Spermophilus dauricus* and find that both pulse and press disturbances have distinct and predictable consequences on the population structure. These examples reinforce the value of our framework across life-histories and complements recent work on demographic resilience (Capdevila et al. 2020; Dakos and Kéfi 2022; MacDonald et al. 2024), which emphasizes the role of transient dynamics in buffering populations. By explicitly unifying pulse and press disturbances, we extend these approaches to a broader class of ecological perturbations and underscore the relevance for both theoretical and applied ecologists.

Our analysis leads to a new matrix, transient response matrix (TRM, **J**_0_), which predicts structure shifts following a press disturbance. When a pulse disturbance acts on a structured population, the resulting transient dynamics are characterized by the damping matrix **Q**_1_. From a macroecological perspective, we find further evidence on how demography shapes resilience (Jiang et al. 2022; Capdevila et al. 2020). First, the largest eigenvalue of TRM is positively correlated with generation time (*T*_*c*_). Second, the TRM is in fact essentially equal to the cumulative distance to stability after a pulse, as defined by Cohen (1979a).

The work allows us to unify disturbance impacts in a single algebraic framework. We focus on small disturbances, similar to what is done for a linear sensitivity analyses. However, in matrix population models, a small press produces a linear change in the SSD but a nonlinear change in growth rate (fitness) and thus our approach may be akin to a nonlinear sensitivity analysis. Our approach should help researchers examine impacts of linear and nonlinear selection on vital rates such as survival, growth, and reproduction. The second derivatives of population growth rate have evolutionary implications (Doak et al. 1994; Brodie et al. 1995; Vasseur and Fox 2007; Shyu and Caswell 2014). They reveal: (i) whether the average fitness of individuals in the population changes linearly as vital rates are perturbed (e.g., Caswell (2001)); (ii) if fitness is nonlinear, whether fitness is concave or convex (Kajin et al. 2025)); and (iii) if the second derivative of fitness with respect to vital rates is positive for observed life histories, then the observed value corresponds to a local minimum, and *vice versa* (Brodie et al. 1995). Here, property (iii) may be particularly useful when examining selection for optimal life history strategies (Charlesworth 1994).

We consider structured populations described by a MPM in a constant environment, but the analysis may be generalized to other contexts, such as community composition and nonlinear multi-species interactions (Bender et al. 1984; Collins et al. 2020; Ratajczak et al. 2017). For example, suppose we have a population vector **n** following a nonlinear MPM with matrix **A**(*θ*, **n**) with parameters *θ*, and dynamics

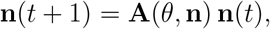

and there is an equilibrium population **n**^*∗*^ that is locally stable. Here of course, vector **n**(*t*) may have components that are stages of one-species or of many species. Now consider a press that changes the parameters to *θ* to (*θ* + *ϵ ϕ*). Then the original stable equilibrium changes to say **n**^*∗*^ + **x**, and our methods can be directly used to compute the change **x**. We plan to follow this direction of analysis in later work.

Our findings are also relevant for stochastic population dynamics and linear response theory (Ruelle 2009; De Nittis and Lein 2017; Jaggi et al. 2024a). In particular, a population in a stochastic environment can be viewed in terms of a sequence of unequal pulses, and their cumulative effects. So for example, we can examine how shifts in mean, variance, and temporal autocorrelation may impact a population’s ability to persist (Drake 2005; Vasseur and Fox 2007). Indeed we note that the long-run stochastic growth rate in a serially correlated environment is given by a quantity that is quite similar to the TRM of this paper, but extended to include the stochastic autocorrelation in the environment (Tuljapurkar and Haridas 2006). We also expect that our methods may be useful in studying the interplay between pulse and press disturbances while incorporating density dependence (Jaggi et al. 2024b) or nonlinear interactions within ecological communities (Medeiros et al. 2023; Medeiros and Saavedra 2023).

Our approach complements earlier work on MPMs. Some general results are found in Cohen’s work on the convexity of the dominant eigenvalue (Cohen 1981; Cohen 1979a; Cohen 1979b; Cohen 1980) used by, e.g., Drake (2005), with more recent work by McCarthy et al. (2008) and Stott (2016). We hope these insights into the connections between nonlinearity, transients, and selection pressures will be useful in future research on understanding and managing population dynamics in a changing world.

## Acknowledgments

RSG was supported by a NERC Pushing the Frontiers grant (NE/X013766/1); MK was supported by the European Commission through the Marie Skłodowska-Curie fellowship (MSCA MaxPersist #101032484) hosted by RSG.

## Appendix

### A. Pulse Disturbances and Transients

#### A.1 Basic Decompositions

We decompose the population matrix as in the main text. Note that

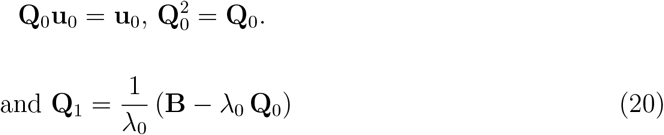

#### A.2 The perpendicular part of population structure following pulse

Suppose a pulse disturbance modifies the matrix **B** to **B** + **D** for one time step at *t* = 0. The matrix **D** is assumed to be zero everywhere except at the entry being perturbed. Since this is a pulse disturbance, the projection matrix reverts to **B** for the next time steps. Here, we assume that the perturbation is small enough so that normalization to total size is not needed. After the pulse, the population structure becomes:

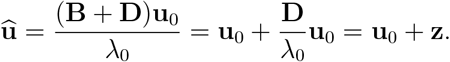

where 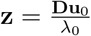 is the shift in population structure caused by the pulse disturbance. However, we are interested in the transient part of the new structure, that is the part that eventually dies out. Such a vector would be orthogonal to SSD. Any part of **z** aligned with **u**_0_ is only a rescaling of all stages and thus does not represent a change in population structure.

Let **z**^*∗*^ be the component of **z** that lies entirely in the transient subspace (i.e., orthogonal to **u**_0_ in the sense of the left eigenvector **v**_0_). To extract the transient component, we want a vector **z**^*∗*^ that is not aligned with stable component **Q**_0_. We achieve this by subtracting from **z** its component along **u**_0_. That is-

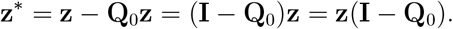

Applying this projection (**I** − **Q**_0_) to **z** yields the orthogonal or transient component:

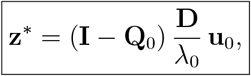

which is Equation (6). Note that the matrix (**I** − **Q**_0_) satisfies:

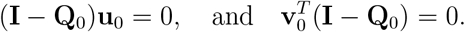

This orthogonality is given by 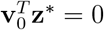, or equivalently **Q**_0_**z**^*∗*^ = 0.

### B Perturbation theory for a Press disturbance

Our analysis follows the perturbation approach in Chapter 2 of Kato (1976). For a press, We examine the effects of changing the matrix **B** by a small amount to **B**+*ϵ***D**. This disturbance in the matrix elements of **B** at time *t*_0_ will produce changes in the dominant eigenvalue and corresponding right eigenvector as,

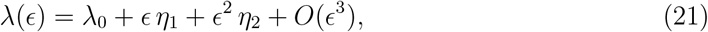

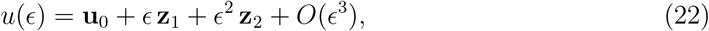

so that

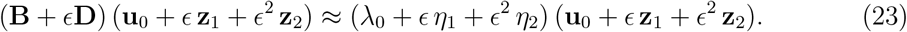

Multiplying through in equation (23) and collecting terms in *ϵ*^0^, *ϵ*^1^, *ϵ*^2^ leads to three equations,

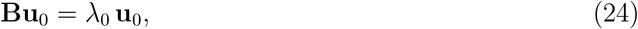

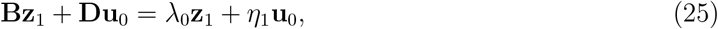

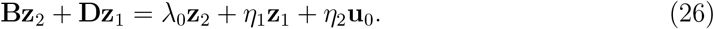

The first of these is obviously true. The others must be used to find the perturbations – the subject of classical perturbation theory. The first-order effects in the middle equation (25) yield the first derivatives of *λ*_0_ and have been applied in ecology Caswell (1996) and Caswell (2001) to study sensitivity and elasticity.

This first-order result is

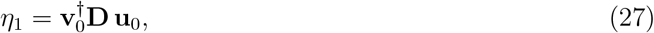

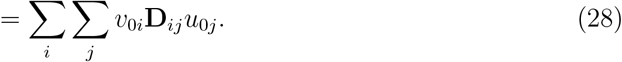

Following Caswell, take the (*i, j*) element of **D** to be 1 (*d*_*ij*_ = 0) and all other elements to be zero, thus yielding the sensitivity

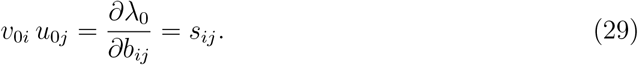

#### B.1 Second Order Perturbation: finding *η*_2_

Second-order perturbations yield *η*_2_. Similar to first-order p erturbation, m ultiply equation (26) by 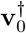 on the left and use orthogonality to see that

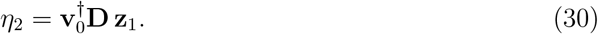

Note that *η*_2_ is a nonlinear response (coefficient of *ϵ*^2^ in equation (26)).

As explained in the references, the first-order analysis along with equation (25) yields

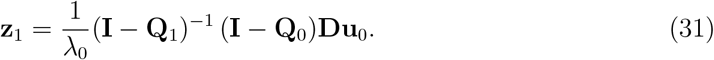

Note that as in the main text, **z**^*∗*^ = (**I** − **Q**_0_)**Du**_0_*/λ*_0_ and so,

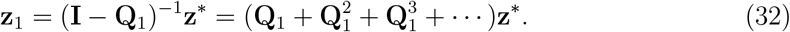

Note that in the main text **Z** = (**I** − **Q**_1_)^−1^*z*^*∗*^ = **z**_1_.

Substitute the value of **z**_1_ to get

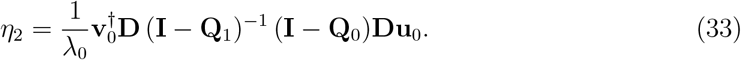

Now define transient response matrix (TRM)

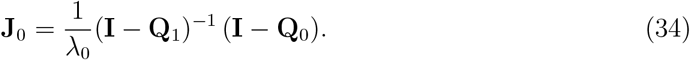

So finally, the second order coefficient is

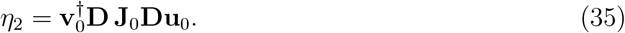

#### B.2 Second order derivatives

Write the change in the dominant eigenvalue as a Taylor series to order *ϵ*^2^,

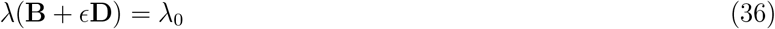

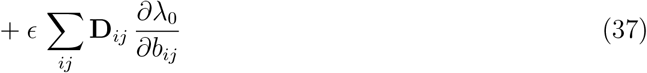

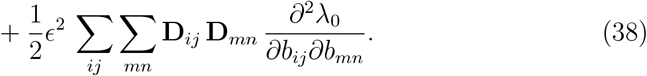

Using this expansion again to order *ϵ*^2^,

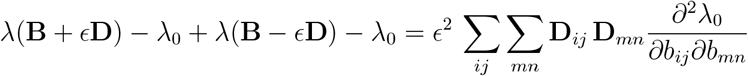

Now use equation (21) and equation (35) to get

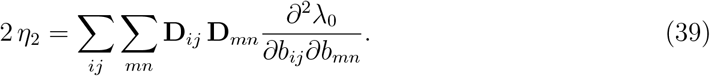

Perturbing only the (*pq*) element yields

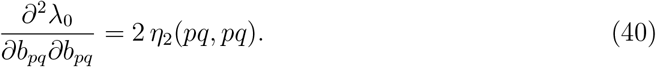

Now perturb distinct elements *p, q* and *k, l*, to find

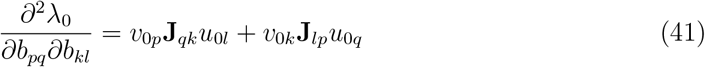

With the sensitivities, *s*_*pl*_ = *v*_0*p*_ *u*_0*l*_ we can write this using Kronecker products as

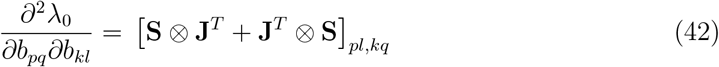

##### Getting to the Hessian - the matrix of second derivatives

Here we construct a matrix, Hessian (**H**), to place all the second derivatives calculated from equation (41). Suppose the population matrix **B** is *S × S*. Then define indices *x, y* that take values 1, 2, …, *S*^2^. We list the elements of **B** columnwise which gives a vector **b**, so *b*_*pq*_ ≡ **b**_*x*_ with

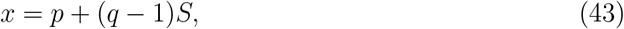

Therefore,

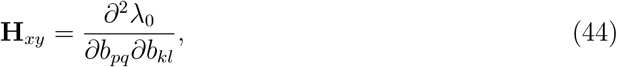

where *x* = *p* + (*q* − 1)*S* and *y* = *k* + (*l* − 1)*S*.

#### B.3 The alternative route to calculate the second derivatives

The second route, I to III and then III to IV. At I the SSD is **u**_0_ and the stable reproductive value is **v**_0_, so in going from I to III the fitness changes by

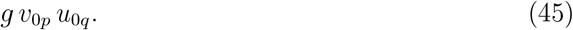

Now we are at III, but there the SSD is to (**u**_0_ + **Z**_2_) with

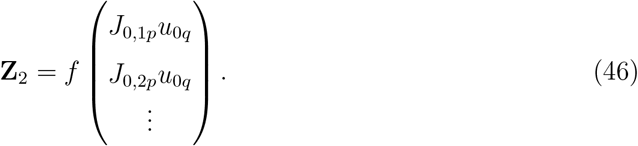

Also at III, the stable reproductive value is now (**v**_0_ + **Y**_2_) with

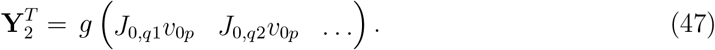

So in the transition III to IV, the fitness changes by the product of

a. the stable proportion in stage *l*, which equation (46) shows is (**u**_0*l*_ + **Z**_2*l*_),
b. the change in the rate, *f*,
c. the stable reproductive value in stage *k*, which equation (47) shows is (**v**_0*k*_ + **Y**_2*k*_).

Thus the total change in the presses from I to III, and III to IV (see Figure 3) is

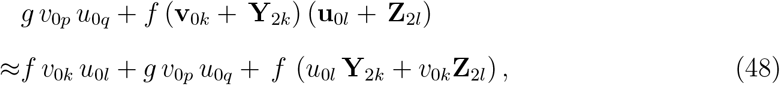

where the higher order term *g***Y**_2*k*_**Z**_2*l*_ is ignored because it is close to zero. Split this up into two bits,

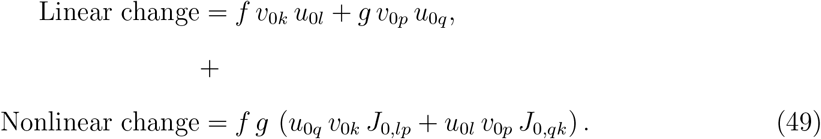

Split this up into two bits,

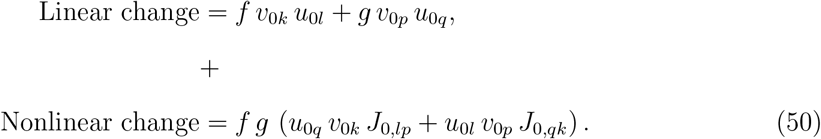

The nonlinear change yields the second derivatives of fitness

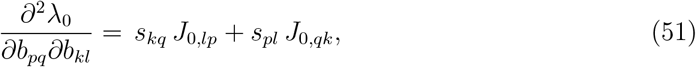

#### B.4 Stepwise Calculation of Second derivatives for *Phaseolus lu-natus*

The matrix for *Phaseolus lunatus* is

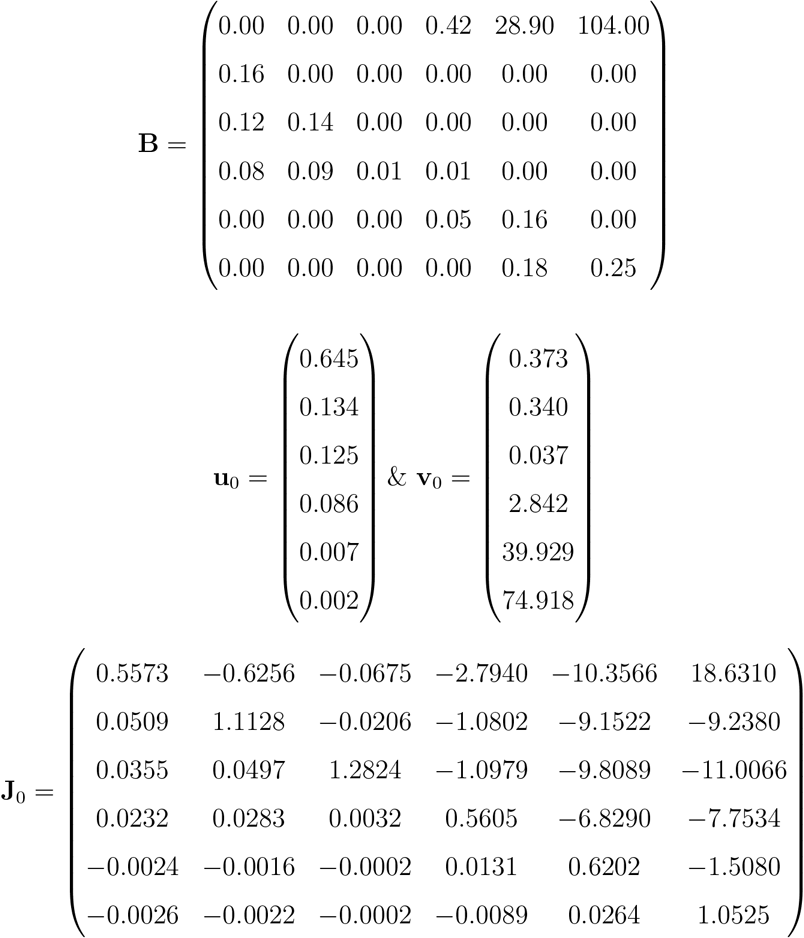

To obtain the second derivative of growth rate, we need the marginal effect of perturbing two demographic rates. For ease of exposition, say we make a press disturbance and change (2% of the element’s value) the (1, 5) and (6, 5) matrix elements to *b*_15_ + 0.578 and *b*_65_ + 0.0036, where the change *g* = 0.578 and *f* = 0.0036.

Fig 3 illustrates two distinct ways of carrying out the above press disturbance, which must lead to the same overall change in fitness.

Say we use the first route, A to B and then B to D. In the change from A to B, our simple argument above shows that the change in fitness is the product

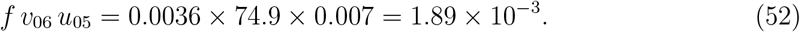

Now we want to make the change from B to D. But at B, the SSD has already changed to (**u**_0_ + *f* **Z**_1_) with

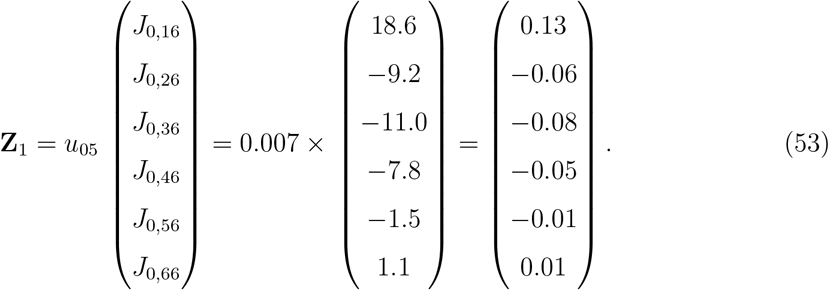

Also at B, the stable reproductive value has also already changed to (**v**_0_ + *f* **Y**_1_) with

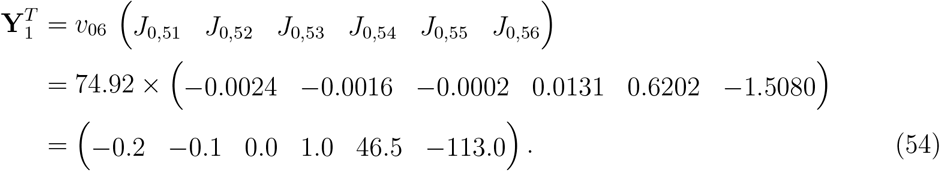

Now we are ready to use our approach to say that in the transition B to D, the fitness changes by the product of

a. the stable proportion in stage 5, which equation (53) shows is (*u*_05_ + *f Z*_15_) = 7.02 *×* 10^−3^,
b. the change in the rate, *g* = 0.578,
c. the stable reproductive value in stage 1, which equation (54) shows is (*v*_01_ + *f Y*_11_) = 0.372.

The product of these terms has to be added to the change (equation (52)) to get the total change in growth rate A to B to D,

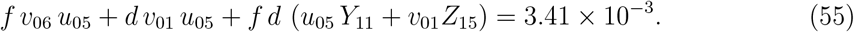

Using (53 – 54) the total change is the sum

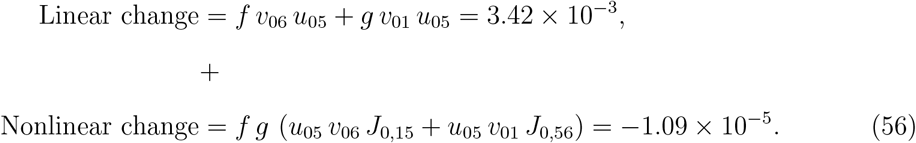

As shown in Fig 3, we could alternatively go from A to C and then C to D. That process involves distinct changes to the SSD and reproductive value. But we get the same final result as in equation (56).

We conclude that the nonlinearity is revealed by making two press disturbances. Think about these changes in terms of the second derivatives of fitness to find

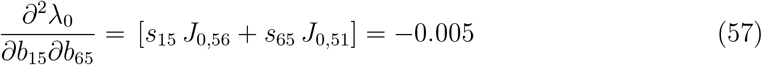

where we have used the sensitivities equation (12). Note that our expression for the second derivative is symmetric with respect to an exchange of the elements *b*_15_, *b*_65_ (as it should be).

The curvature of fitness as measured by the second derivatives in equation (57) depends on TRM (**J**_0_). Consequently any analysis of second derivatives will provide detailed information about TRM. The next section describes the many connections between TRM and transient dynamics.

### C When the population matrix has distinct eigenvalues

Write 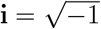, and the higher eigenvalues as

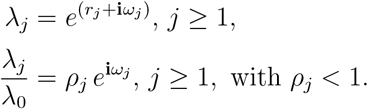

The ratios *ρ*_*j*_ = *λ*_*j*_*/λ*_0_ *<* 1 are called the damping ratios. Each eigenvalue has a corresponding right, left eigenvector, 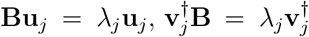, and we set 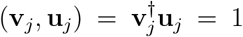 (the subscript *†* indicates the combination of a transpose and a complex conjugate). Then we can write a spectral decomposition (Good 1969) of the matrix **B** as

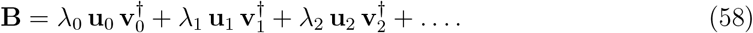

Recalling the definition of **Q**_1_ in (2), we find

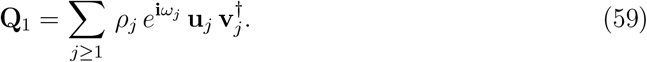

Use a geometric series expansion to find the explicit expression

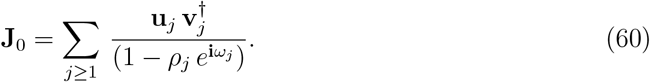

This happens to be a spectral decomposition of **J**_0_, excluding the eigenvector **u**_0_ for which the eigenvalue is 0. Thus for *j* = 1, 2, … the vectors **u**_*j*_ **v**_*j*_ are right, left eigenvectors of **J**_0_ corresponding to eigenvalue 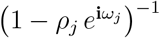.

### D Relation between TRM and Cohen’s cumulative distance

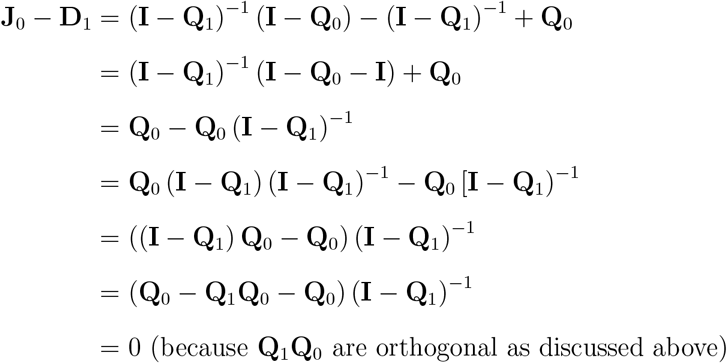

## References

Abbott, Karen C et al. (2024). “When and why ecological systems respond to the rate rather than the magnitude of environmental changes”. Biological Conservation 292, p. 110494.

Amor, DR, C Ratzke, and J Gore (2020). Transient invaders can induce shifts between alternative stable states of microbial communities. Sci Adv 6: eaay8676.

Arnoldi, J-F, Azenor Bideault, Michel Loreau, and Bart Haegeman (2018). “How ecosystems recover from pulse perturbations: A theory of short-to long-term responses”. Journal of theoretical biology 436, pp. 79–92.

Barnosky, Anthony D et al. (2012). “Approaching a state shift in Earth’s biosphere”. Nature 486.7401, pp. 52–58.

Bender, Edward A, Ted J Case, and Michael E Gilpin (1984). “Perturbation experiments in community ecology: theory and practice”. Ecology 65.1, pp. 1–13.

Brodie, Edmund D, Allen J Moore, and Fredric J Janzen (1995). “Visualizing and quantifying natural selection”. Trends in Ecology & Evolution 10.8, pp. 313–318.

Capdevila, Pol, Iain Stott, Maria Beger, and Roberto Salguero-Gómez (2020). “Towards a comparative framework of demographic resilience”. Trends in Ecology & Evolution 35.9, pp. 776–786.

Capdevila, Pol et al. (2021). Reconciling resilience across ecological systems, species and subdisciplines.

Caswell, H. (2001). Matrix population models: construction, analysis and interpretation. 2nd. Sunderland, Mass.: Sinauer associates, Sunderland, Mass.

Caswell, Hal (1996). “Second derivatives of population growth rate: calculation and applications”. Ecology, pp. 870–879.

Charlesworth, B. (1994). Evolution in Age-Structured Populations. Cambridge University Press, Cambridge.

Cohen, J.E. (1977). “Ergodicity of age structure in populations with Markovian vital rates, III: finite-state moments and growth rate; an illustration”. Advances in Applied Probability, pp. 462–475.

Cohen, J.E. (1979a). “The cumulative distance from an observed to a stable age structure”. SIAM Journal on Applied Mathematics 36.1, pp. 169–175.

Cohen, J.E. (1980). “Convexity properties of products of random nonnegative matrices”. Proceedings of the National Academy of Sciences 77.7, pp. 3749–3752.

Cohen, Joel E (1981). “Convexity of the dominant eigenvalue of an essentially nonnegative matrix”. Proceedings of the American Mathematical Society 81.4, pp. 657–658.

Cohen, J.E. (1979b). “Random evolutions and the spectral radius of a non-negative matrix”. Mathematical Proceedings of the Cambridge Philosophical Society 86.2, pp. 345–350. doi: 10.1017/S0305004100056164.

Collins, Scott L et al. (2020). “Press–pulse interactions and long-term community dynamics in a Chihuahuan Desert grassland”. Journal of Vegetation Science 31.5, pp. 722–732.

Dakos, Vasilis and Sonia Kéfi (2022). “Ecological resilience: what to measure and how”. Environmental Research Letters 17.4, p. 043003.

De Nittis, Giuseppe and Max Lein (2017). Linear response theory: an analytic-algebraic approach. Springer.

Doak, Daniel, Peter Kareiva, and Brad Klepetka (1994). “Modeling population viability for the desert tortoise in the western Mojave Desert”. Ecological Applications 4.3, pp. 446– 460.

Donohue, Ian et al. (2016). “Navigating the complexity of ecological stability”. Ecology letters 19.9, pp. 1172–1185.

Drake, J.M. (2005). “Population effects of increased climate variation”. Proceedings of The Royal Society B 272.1574, p. 1823.

Gaillard, J.M. et al. (2005). “Generation Time: A Reliable Metric to Measure Life-History Variation among Mammalian Populations”. American Naturalist 166.1, pp. 119–123.

Glasby, Tim M and AJ Underwood (1996). “Sampling to differentiate between pulse and press perturbations”. Environmental monitoring and assessment 42, pp. 241–252.

Good, Irving John (1969). “Some applications of the singular decomposition of a matrix”. Technometrics 11.4, pp. 823–831.

Haridas, CV and Shripad Tuljapurkar (2007). “Time, transients and elasticity”. Ecology Letters 10.12, pp. 1143–1153.

Hastings, Alan (2001). “Transient dynamics and persistence of ecological systems”. Ecology Letters 4.3, pp. 215–220.

IBPGR (1982). International Board for Plant Genetic Resources (1982) Lima bean descriptors. Rome (Italy): IBPGR 36 p.

Inamine, Hidetoshi et al. (2022). “Pulse and Press Disturbances Have Different Effects on Transient Community Dynamics”. The American Naturalist 200.4, pp. 571–583.

Jaggi, Harman, David Steinsaltz, and Shripad Tuljapurkar (2024a). “Temporal variability can promote migration between habitats”. Theoretical Population Biology 158, pp. 195–205.

Jaggi, Harman et al. (2024b). “Density dependence shapes life-history trade-offs in a foodlimited population”. Ecology Letters 27.11, e14551.

Jentsch, Anke and Peter White (2019). “A theory of pulse dynamics and disturbance in ecology”. Ecology 100.7, e02734.

Jiang, Sha et al. (2022). “Reproductive dispersion and damping time scale with life-history speed”. Ecology Letters 25.9, pp. 1999–2008.

Kajin, M et al. (2025). “Second-order elasticities for Ecology and Evolution: Unravelling nonlinear fitness responses to perturbations”. bioRxiv, pp. 2025–04.

Kato, Tosio (1976). Perturbation theory for linear operators. Vol. 132. Springer Science & Business Media.

Koons, David N., Todd W. Arnold, and Michael Schaub (2017). “Understanding the demographic drivers of realized population growth rates”. Ecological Applications 27.7, pp. 2102–2115. doi: 10.1002/eap.1594. eprint: https://esajournals.onlinelibrary.wiley.com/doi/pdf/10.1002/eap.1594. URL: https://esajournals.onlinelibrary.wiley.com/doi/abs/10.1002/eap.1594.

Koons, David N., James B. Grand, Bertram Zinner, and Robert F. Rockwell (2005). “Transient population dynamics: Relations to life history and initial population state”. Ecological Modelling 185.2, pp. 283–297. ISSN: 0304-3800. doi: 10.1016/j.ecolmodel.2004.12.011. URL: https://www.sciencedirect.com/science/article/pii/S0304380005000025.

MacDonald, Jane Shaw, Frithjof Lutscher, and Yves Bourgault (2024). “Climate change fluctuations can increase population abundance and range size”. Ecology Letters 27.6, e14453.

McCarthy, Dominic, Stuart Townley, and Dave Hodgson (2008). “On second order sensitivity for stage-based population projection matrix models”. Theoretical Population Biology 74.1, pp. 68–73.

Medeiros, Lucas P, Michael G Neubert, Heidi M Sosik, and Stephan B Munch (2025). “A nonequilibrium framework for community responses to pulse perturbations”. bioRxiv, pp. 2025–05.

Medeiros, Lucas P and Serguei Saavedra (2023). “Understanding the state-dependent impact of species correlated responses on community sensitivity to perturbations”. Ecology 104.8, e4115.

Medeiros, Lucas P et al. (2023). “Ranking species based on sensitivity to perturbations under non-equilibrium community dynamics”. Ecology Letters 26.1, pp. 170–183.

Morozov, Andrew et al. (2020). “Long transients in ecology: Theory and applications”. Physics of life reviews 32, pp. 1–40.

Morozov, Andrew et al. (2024). “Long-living transients in ecological models: Recent progress, new challenges, and open questions”. Physics of Life Reviews.

Neubert, Michael G, Hal Caswell, and JD Murray (2002). “Transient dynamics and pattern formation: reactivity is necessary for Turing instabilities”. Mathematical biosciences 175.1, pp. 1–11.

Ratajczak, Zak et al. (2017). “The interactive effects of press/pulse intensity and duration on regime shifts at multiple scales”. Ecological Monographs 87.2, pp. 198–218.

Refsland, Tyler and Jennifer Fraterrigo (2018). “Fire increases drought vulnerability of Quercus alba juveniles by altering forest microclimate and nitrogen availability”. Functional Ecology 32.10, pp. 2298–2309.

Ripple, William J and Robert L Beschta (2012). “Trophic cascades in Yellowstone: the first 15 years after wolf reintroduction”. Biological Conservation 145.1, pp. 205–213.

Ruelle, David (2009). “A review of linear response theory for general differentiable dynamical systems”. Nonlinearity 22.4, p. 855.

Salguero-Gómez, Roberto et al. (2015). “The compadre P lant M atrix D atabase: an open online repository for plant demography”. Journal of Ecology 103.1, pp. 202–218.

Salguero-Gómez, Roberto et al. (2016). “COMADRE: a global data base of animal demography”. Journal of Animal Ecology 85.2, pp. 371–384.

Shyu, Esther and Hal Caswell (2014). “Calculating second derivatives of population growth rates for ecology and evolution”. Methods in Ecology and Evolution 5.5, pp. 473–482.

Sironi, Maria (2019). “Fertility histories and chronic conditions later in life in Europe”. European journal of ageing 16.3, pp. 259–272.

Stott, Iain (2016). “Perturbation analysis of transient population dynamics using matrix projection models”. Methods in Ecology and Evolution 7.6, pp. 666–678.

Tao, Yun et al. (2021). “Transient disease dynamics across ecological scales”. Theoretical Ecology 14.4, pp. 625–640.

Traill, Lochran W, Susanne Schindler, and Tim Coulson (2014). “Demography, not inheritance, drives phenotypic change in hunted bighorn sheep”. Proceedings of the National Academy of Sciences 111.36, pp. 13223–13228.

Tuljapurkar, S. and CV Haridas (2006). “Temporal autocorrelation and stochastic population growth”. Ecology Letters 9.3, pp. 327–337.

Vasseur, David A and Jeremy W Fox (2007). “Environmental fluctuations can stabilize food web dynamics by increasing synchrony”. Ecology Letters 10.11, pp. 1066–1074.

White, J Wilson et al. (2013). “Transient responses of fished populations to marine reserve establishment”. Conservation Letters 6.3, pp. 180–191.

Williams, Jennifer L et al. (2011). “Distance to stable stage distribution in plant populations and implications for near-term population projections”. Journal of ecology 99.5, pp. 1171–1178.

Yang, Louie H, Justin L Bastow, Kenneth O Spence, and Amber N Wright (2008). “What can we learn from resource pulses”. Ecology 89.3, pp. 621–634.

